# Synchronized path-integration recalibration but distinct landmark-control dynamics in head direction and CA1 place cells

**DOI:** 10.1101/2025.10.08.681135

**Authors:** Ravikrishnan P. Jayakumar, Yotaro Sueoka, Marissa Ferreyros, Brian Y. Li, Manu S. Madhav, Xingyu Chen, James J. Knierim, Noah J. Cowan

**Author notes:** **Corresponding authors:** Ravikrishnan P. Jayakumar, Yotaro Sueoka, James J. Knierim, Noah J. Cowan. These authors contributed equally (Ravikrishnan P. Jayakumar, Yotaro Sueoka). These authors jointly supervised this work (James J. Knierim, Noah J. Cowan).

## Abstract

Accurate spatial navigation relies on path integration, a process of tracking one’s location by integrating self-motion cues. Path integration uses a gain factor relating self-motion signals to displacement on the cognitive map. This gain is plastic, recalibrating rapidly to match perceived displacements relative to external cues. To elucidate the mechanism of recalibration, we simultaneously recorded from place cells, which instantiate the cognitive map, and head direction (HD) cells, thought to orient the map. Persistent conflict between self-motion and visual feedback induced functionally identical recalibration of path-integration gain in the two neural populations during forward locomotion; however, during locomotor immobility accompanied by head-scanning, HD cells did not exhibit recalibration. Moreover, the two populations manifested differential field-shifting dynamics relative to landmarks during recalibration. These results uncover a tightly coordinated yet behavior-dependent recalibration process across the navigation circuit that achieves a robust yet flexible coupling of the internal sense of position and direction.

## Introduction

From foraging ants to migrating birds (1–3), animals display remarkable navigational abilities. In some animals, this capacity for spatial navigation is thought to rely in part on a “cognitive map” (4–6)—a neural representation of the surrounding environment. For successful navigation, an animal must know its location within this cognitive map and continuously update its position as it moves through space. While salient external landmarks (such as the sun’s position, distant mountains, or a familiar building) can help pinpoint location and orientation (7,8), animals also possess an internal capacity to navigate using signals generated by self-motion. This internal computation, called path integration, relies on integrating self-motion signals, including vestibular input, motor efference copy, and proprioception, to estimate linear and angular displacement (9–17).

Because of the inherent noise in the biological signals being integrated, the position estimated by path integration can accumulate errors. External cues, when available, can correct such noise-driven errors (18). However, a consistent bias in these errors (i.e., consistently under-or over-estimating distance traveled relative to landmarks) can indicate to the brain that the path integration computation itself may be miscalibrated. Human behavioral studies have demonstrated that the miscalibrated computation corrects itself to match perceived displacements relative to external cues (19,20). Our previous research, recording from the rodent hippocampus—the proposed seat of the cognitive map (4)—demonstrated rapid recalibration of internal path integration computation in response to such biases (21,22). This recalibration involves adjusting the path-integration gain—the factor converting self-motion into expected displacement. Specifically, when animals experienced a mismatch between their self-motion and visual feedback, indicating they were moving faster or slower than their internal estimate, their path-integration gain adapted to rescale the integrated self-motion to better match the visually perceived speed and displacement. While this plasticity was observed in hippocampal recordings, it is not known how recalibration is coordinated across the brain regions involved in spatial navigation.

Given that the animals in our prior studies ran laps around a circular track, biased errors were introduced into both linear *and* angular path integration computations. It is unclear how recalibration would be expressed in these components of the path integration computation. To address this, we focus on two cell types for this study: hippocampal place cells and head direction cells. Place cells fire in specific locations within an environment (23) and are influenced by both linear and angular self-motion. Head direction (HD) cells fire selectively when the animal’s head points in a specific direction (24,25) and are primarily driven by angular self-motion (26–30). These two cell types are strongly interconnected: HD cells are believed to set the orientation of the place cell map (31–35), while place cells may provide feedback to the HD system (36–38). While place cells and HD cells generally maintain coherent activity (31,39–41), they can dissociate when presented with conflicting external reference frames (41). In addition, place cells and HD cells display different patterns of backward shifting in familiar and cue-conflict environments (42), a phenomenon that is thought to reflect Hebbian plasticity (43) or behavioral time-scale dependent plasticity (44) mechanisms that store learned sequences in memory. Such dissociations between the place and HD cell representations might also occur in the event of differential recalibration of the linear and angular components of path integration, providing differential path-integration drive onto place cells and HD cells during forward movement. Furthermore, the extent of recalibration in the angular path integration system could be independently assessed by examining HD cell activity during periods of immobility and head-scanning, when there is no linear movement.

To address these questions, we simultaneously recorded from hippocampal CA1 place cells and HD cells, mostly from the thalamus (45–47) and a small minority from the cingulum bundle (48,49) of rats. Rats navigated in a circular virtual-reality (VR) apparatus, the Dome (50), that allows precise control over the relationship between non-visual self-motion cues and visual landmark movement, creating a conflict between the internal and external estimates of displacement. We found that during locomotion, place cells and HD cells undergo tightly coupled recalibration, exhibiting functionally identical recalibrated path-integration gains. However, during periods of immobility and head-scanning, HD cells displayed an approximately unitary gain, suggesting that the angular velocity input to the HD cells did not experience a gain change under these conditions or, alternatively, that the expression of path-integration gain in the HD cell system may depend on behavioral context.

## Results

To study the coordination between hippocampal and head direction (HD) systems during path integration recalibration, CA1 place cells and HD cells from multiple regions (**Figure S1a**) were simultaneously recorded from 4 rats (1 male, 3 female, 39 sessions; mean 17.6 CA1 units/session, 2 HD units/session meeting place-cell and HD-cell inclusion criteria, see Methods) in the Dome VR system (Figure 1a, see Methods). Six sessions did not have simultaneously recorded HD cells and are used only in CA1-specific analyses. Animals ran laps around the periphery of a circular table (lap circumference = 4.2 m) positioned under a hemispherical shell. A constellation of salient 2D white visual cues was projected onto the inner surface of the shell. The projected cues were moved azimuthally as a linear function of the animal’s movement. The relationship between visual landmark movement and the animal’s movement was defined by an experimental gain, *G* (**Figure 1b**). When *G* = 1, the landmarks were stationary. When *G* > 1, landmarks moved in the direction opposite to the rat’s movement, creating the illusion of faster motion. When *G* < 1, landmarks moved in the same direction as the rat, creating the illusion of slower motion. A typical experimental session consisted of four epochs (**Figure 1c**). In Epoch 1, the landmarks were stationary (*G* = 1). In Epoch 2, *G* was gradually increased or decreased to a final value, *G_final_*. In Epoch 3, *G* was held constant at *G_final_*. Epochs 1-3 were designed to gradually transition from the animal’s everyday learned relationship between self-motion and stationary landmarks to a sustained period of an altered relationship— scaled by *G_final_*—between landmarks and non-visual self-motion cues. Finally, in Epoch 4, the visual landmarks were removed, leaving only a dim, circularly symmetric ring of illumination near the top of the dome. In the absence of salient external cues, the neural updates of position and direction during this epoch were presumably driven primarily by path integration, allowing us to measure any persistent adaptation or recalibration of the path-integration gain resulting from the cue conflict introduced in the preceding epochs, as done in prior studies (21,22,51).

**Figure 1:**
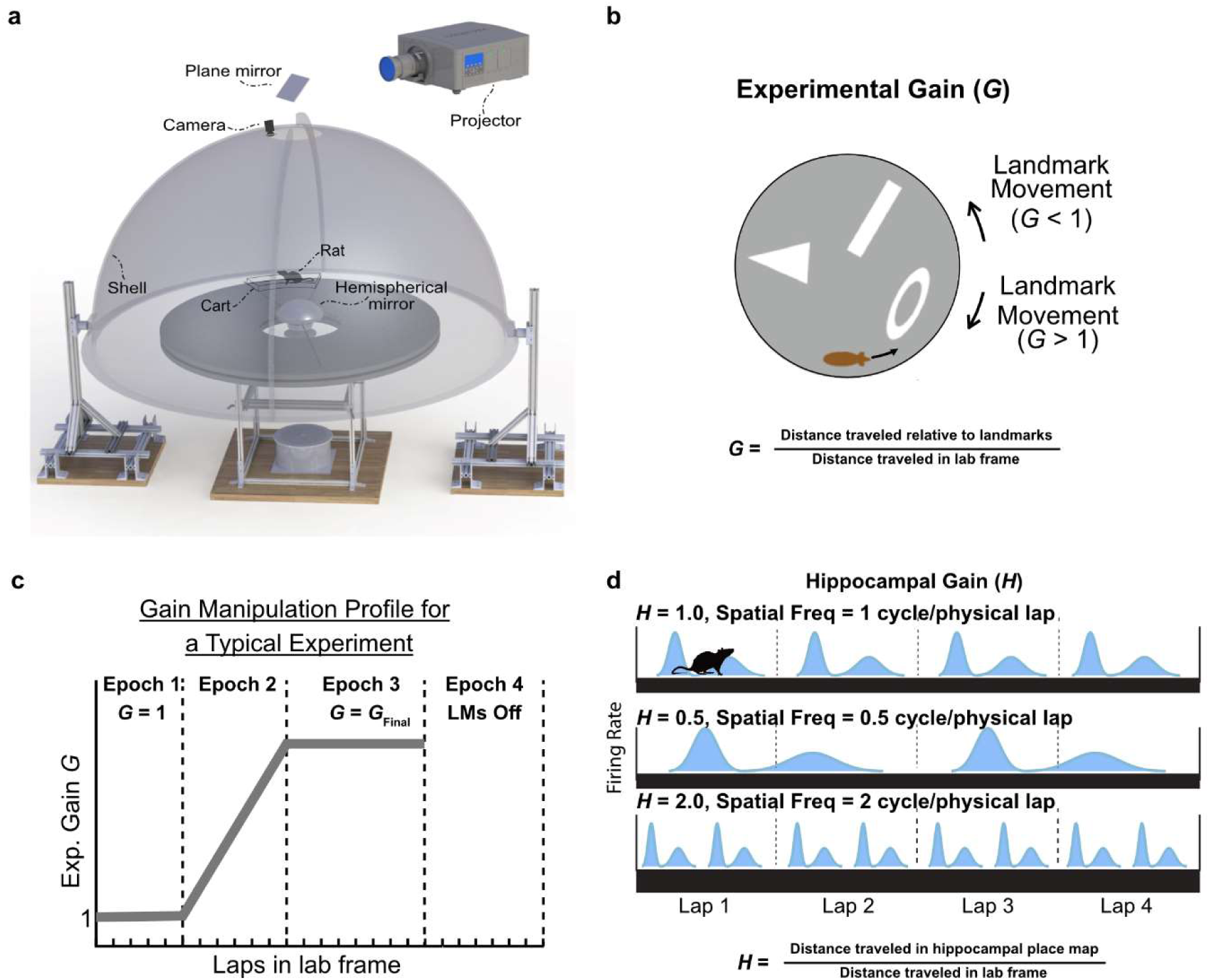
Experiment apparatus, visual cue gain manipulation protocol, and measurement of hippocampal gain. **(a)** Schematic of the Dome virtual-reality (VR) system. A rat locomotes on a circular table located beneath a hemispherical projection shell. Visual landmarks are projected onto the inner surface of the dome via a mirror system. A boom arm, attached to a central rotating pillar, either directly harnesses the animal or supports a carriage that follows the animal’s movement (see Methods for details). **(b)** Illustration of experimental gain manipulation. Visual landmarks inside the Dome move in relation to the rat’s movement, controlled by the gain *G*, defined as the ratio of the rat’s movement in landmark frame to its movement in lab frame. *G* > 1 indicates perceived movement greater than actual movement, while *G* < 1 indicates perceived movement less than actual movement. **(c)** Gain manipulation profile across a typical experiment, divided into Epochs 1-4. Experimental gain (Exp. G) is manipulated in Epochs 2 to 3 and set to the final gain value (*G_final_*) in Epoch 3. Landmarks are removed in Epoch 4 (LMs Off). **(d)** Illustration of place cell firing rate maps, depicting the relationship between hippocampal gain (*H*) and spatial firing frequency. Examples show firing rate maps for *H* = 1 (one field per lap), *H* = 0.5 (one field every two laps), and *H* = 2 (two fields per lap).

When a place cell is anchored by visual landmarks, its firing pattern repeats once per *virtual* lap (i.e., a lap in the landmark reference frame). In the case of *G* = 1 (stationary landmarks), the firing pattern of a place cell also repeats once per physical lap. However, as the gain is manipulated, the spatial frequency of firing changes with respect to the physical track. For example, at *G* = 2, the firing pattern repeats every *half of* a physical lap (or two times per physical lap, maintaining a constant firing location relative to the landmarks). This spatial frequency of place cell firing, defined as the cell’s gain (*H_i_*), represents the readout of the animal’s rate of travel through its place map relative to its rate of physical travel (**Figure 1d**). In most sessions, the firing of the place cell ensemble remained anchored to the landmarks (**Figure S1b**), i.e. *H_i_* = *G*. The gains estimated from individual CA1 cells fell into a tight distribution, indicating a coherent ensemble response (**Figure S1c-d**), as expected from previous work (21,22). Given the coherent response of the CA1 population (except in 2 sessions, discussed in the next Results section), we assume a single hippocampal gain, *H*, calculated as the median of the continuous gain traces *H_i_* of simultaneously recorded place cells in a session. Sessions where the place cell ensemble remained anchored to the landmark cues were classified as “landmark-control”; sessions where anchoring was absent or lost for prolonged periods of the session were classified as “landmark-failure” (see **Methods; Figure S2**). Place cell and HD cell responses during forward running were directly compared by defining the gain of HD cells as the angular frequency of firing of the HD cell, similar to spatial frequency for place cells. The gains of simultaneously recorded head direction cells, *D_i_*, responded as a coherent population in sessions with simultaneously recorded units from more than one region (**Figure S1e**; (52)). Given the coherent HD response, we treated the median HD cell gain, *D,* as the gain of the HD system.

### HD cells and place cells were tightly coupled in both landmark-control and landmark-failure sessions

Place fields and HD cell tuning curves were anchored to the landmarks in most sessions (26/33 sessions). **Figure 2a** shows an example landmark-control session. During gain manipulation (Epochs 2-3), simultaneously recorded place and HD cells consistently fired at a stable location and head yaw angle, respectively, in the landmark reference frame but appeared untuned in the lab reference frame; this is reflected by the fact that the hippocampal gain and HD gain followed the experimental gain (**Figure 2a, top panel**). During forward locomotion in Epoch 3 (*G = G_final_ ≠* 1), HD cells in landmark-control sessions showed poor directional tuning in the laboratory frame (**Figure 2b**) but maintained their tuning in the landmark frame (**Figure 2c**). The median hippocampal gain and HD gain for each landmark-control session closely matched the experimental gain (**Figure 2d; Figure S3, landmark-control panels**). These results confirm that, under conditions of landmark control, HD cells remained anchored to the landmarks along with the place cell population, consistent with previous findings (31,53,54).

**Figure 2:**
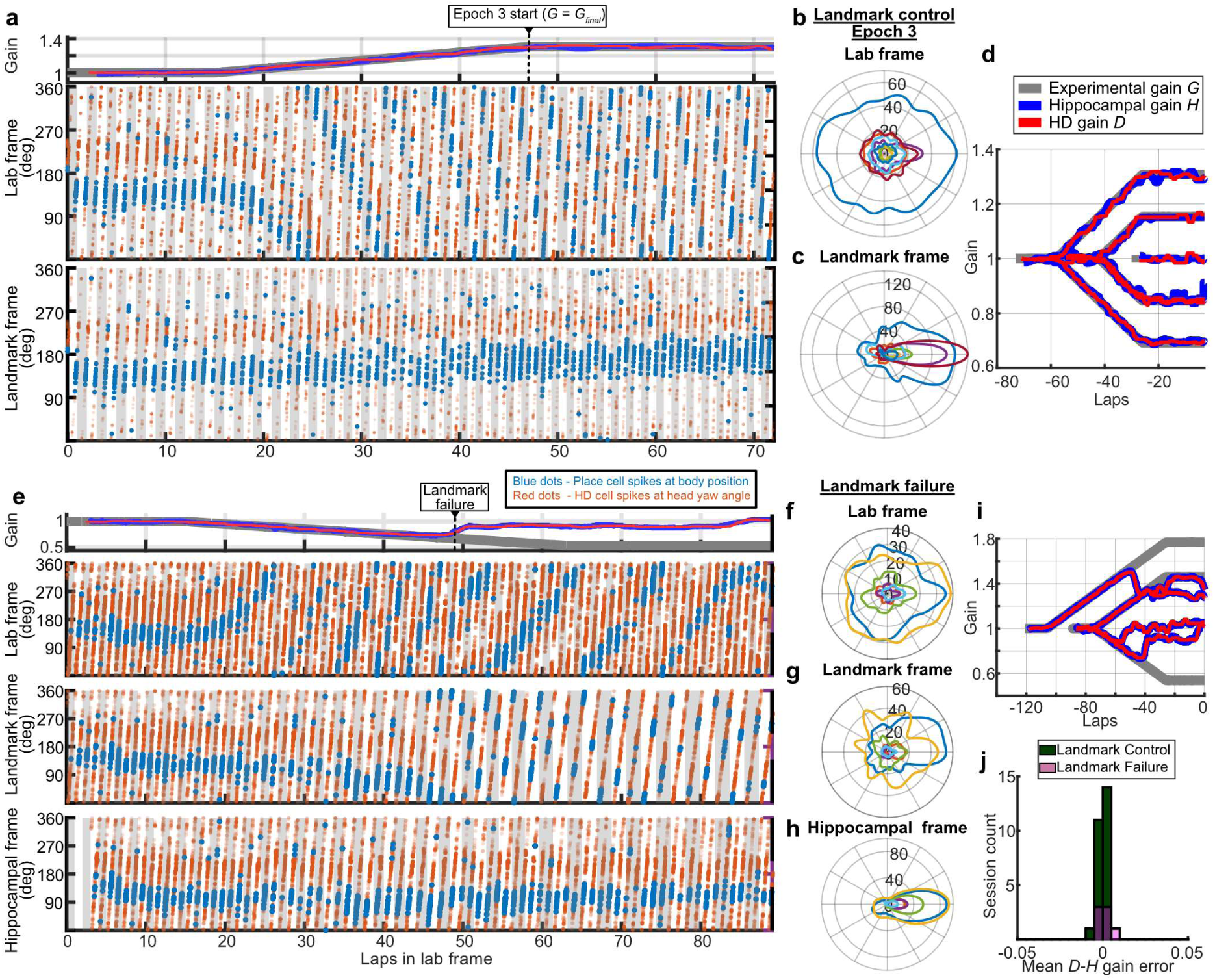
CA1 place cells and HD cells exhibit tightly coordinated gains under landmark manipulations in VR. **(a)** Example raster plot of a place cell and a simultaneously recorded HD cell from a landmark-control session for Rat 1078. The x-axis represents the cumulative distance traveled by the rat in the lab frame. *Top panel:* The hippocampal population gain (blue line), HD gain (red line), and the experimental gain (gray line, frequently hidden behind red and blue lines) for a landmark-control session. The tight correspondence between these 3 gains indicates that the firing fields of the place cells and HD cells were strongly controlled by the rotating landmark frame of reference. *Middle panel:* Body angular position at the instant of place cell spikes (blue dots) and head angle at HD cell spikes (red dots) in the lab frame (y-axis) during forward locomotion. Place cell and HD cell firing fields changed location and direction as the experimental gain ramped from 1 to 1.31. *Bottom panel:* The same spikes as in the middle panel, replotted with respect to the landmark frame (y-axis). The consistent horizontal alignment of spikes in the landmark frame indicates that the cells’ spatial and directional tuning were stable relative to the landmarks, not the lab frame. In both middle and bottom panels, alternating gray and white bars delineate individual laps around the track in lab and landmark frames, respectively. **(b)** HD tuning curves during forward locomotion (forward speed > 5.8 cm/s) for all HD cells recorded from Rat 1078 across Epoch 3 of all landmark-control sessions, plotted in the lab frame, show disrupted directional tuning. **(c)** Firing rates of the same HD cells as in (b) plotted in the landmark frame show sharp directional tuning. **(d)** Gain traces from all the landmark-control sessions from Rat 1078 (see Figure S3 for all rats). Traces show experimental gain *G* (gray), median hippocampal population gain *H* (blue), and median HD gain *D* (red). The gain of the HD cells closely tracks both the experimental gain and the hippocampal gain.

While the place cell map remained anchored to the landmark frame of reference in most sessions, in a subset of sessions (7/33 sessions), the internal map decoupled from the landmarks, presumably caused by errors between idiothetic and allothetic cues introduced due to gain manipulation indicating landmark instability (55). In a raster plot from an example landmark-failure session (**Figure 2e**), an HD cell decoupled from the landmarks at the same time as a simultaneously recorded CA1 place cell, and the HD tuning curve subsequently drifted together with the CA1 place field; as such, HD tuning was better maintained in the *hippocampal frame of reference* (i.e., relative to the activity of the place cell population) compared to the lab and landmark frames (**Figure 2f-h**). Comparison of hippocampal gain and HD gain across all landmark-failure sessions shows that the two populations coherently broke away from the landmarks and drifted together afterwards (**Figure 2i; Figure S3**). Aggregating across all sessions and animals, the HD gain exhibited minimal deviation from the hippocampal gain for both landmark-control and landmark-failure sessions (**Figure 2j**). This tight coupling between the two populations is consistent with a previous study showing that place and HD cells can drift relative to static visual cues before settling into a new stable mapping, with HD and place cells maintaining coherence on fine time scales (31).

There were two anecdotal sessions that violated the strict coupling typically observed between place cells and HD cells (**Figure S4**; these sessions are not included in the session count of 39 nor in other analyses). In these two sessions, a more complex coupling emerged that was dependent on periodic conjunctions of a stable external frame and a rotating HD frame. HD cell tuning curves drifted relative to the landmark/lab frames (which were coincident as *G* = 1) while the place cell population split into two subsets: one subset drifted together with the HD cells and the other subset was active only at a specific conjunction of the landmark/lab frame and the drifting HD frame. Such a conjunctive response has been observed in the retrosplenial cortex (56). In the second of the two sessions, once the visual gain G was increased away from 1, the CA1 subpopulation encoding the conjunction of external and HD reference frames lost its conjunction over the course of a few laps to solely favor, and drift along with, the HD frame.

### HD cells and place cells recalibrated coherently when subjected to sustained conflict between path integration and landmarks

Our previous study (21) showed that the experimental gain applied to the landmarks in Epochs 2-3 caused a recalibration of the internal path-integration gain of the hippocampus. This recalibration was revealed when the landmarks were abruptly turned off in Epoch 4, leaving path integration as the primary driver of the update of the activity of the place cells. *H* in Epoch 4 tended to be greater than its initial value in Epoch 1 when *G_final_* > 1 in Epochs 2-3 and less than its initial value when *G_final_* < 1 in Epochs 2-3. This result demonstrated recalibration of the path-integration gain in a direction that minimized conflict between estimates of location based on path integration and landmark positioning systems. Given the strong coupling of the HD cells and place cells in Epochs 2-3 (**Figure 2**), we next asked how the two populations would recalibrate their path-integration gain to minimize the error between the conflicting landmark and path integration cues. For this analysis, we considered only landmark control sessions and measured the path-integration gain using the spikes from the first six laps (i.e., size of the spatial window used to estimate gain; see Methods) in Epoch 4, as in our prior study.

Figure 3a shows an example recalibration session with a simultaneous recording of a place cell and an HD cell. As in Figure 2, the tuning profiles of both cells drifted in sync in the lab frame when the experimental gain *G* ramped up to its final value of 1.46, but they remained stable in the landmark frame. When the landmarks were removed at the start of Epoch 4, the place cell and HD cell remained tightly coupled with a recalibrated gain of 1.29 (Fig. 3a, top), a value greater than the gain of 1 in Epoch 1. During Epoch 4, HD cells from landmark-control sessions maintained a stable firing direction in the hippocampal frame of reference, reflecting the tight coupling between the two populations (Figure 3b-d). The values of *H* and *D* from all landmark-control sessions from one rat are shown in Figure 3e, illustrating the tight coupling between these values across all epochs, including recalibrated values in Epoch 4. Across all 4 rats in the study, we replicated the linear relationship between experimental gain *G_final_* in Epoch 3 and the recalibrated gain *H_recal_* in Epoch 4 (Figure 3f). The recalibrated gain of HD cells *D_recal_* in Epoch 4 across all sessions precisely matched that of the hippocampus *H_recal_* (Figure 3g**,h**). These findings demonstrate that HD cells recalibrate coherently with hippocampal place cells.

**Figure 3.**
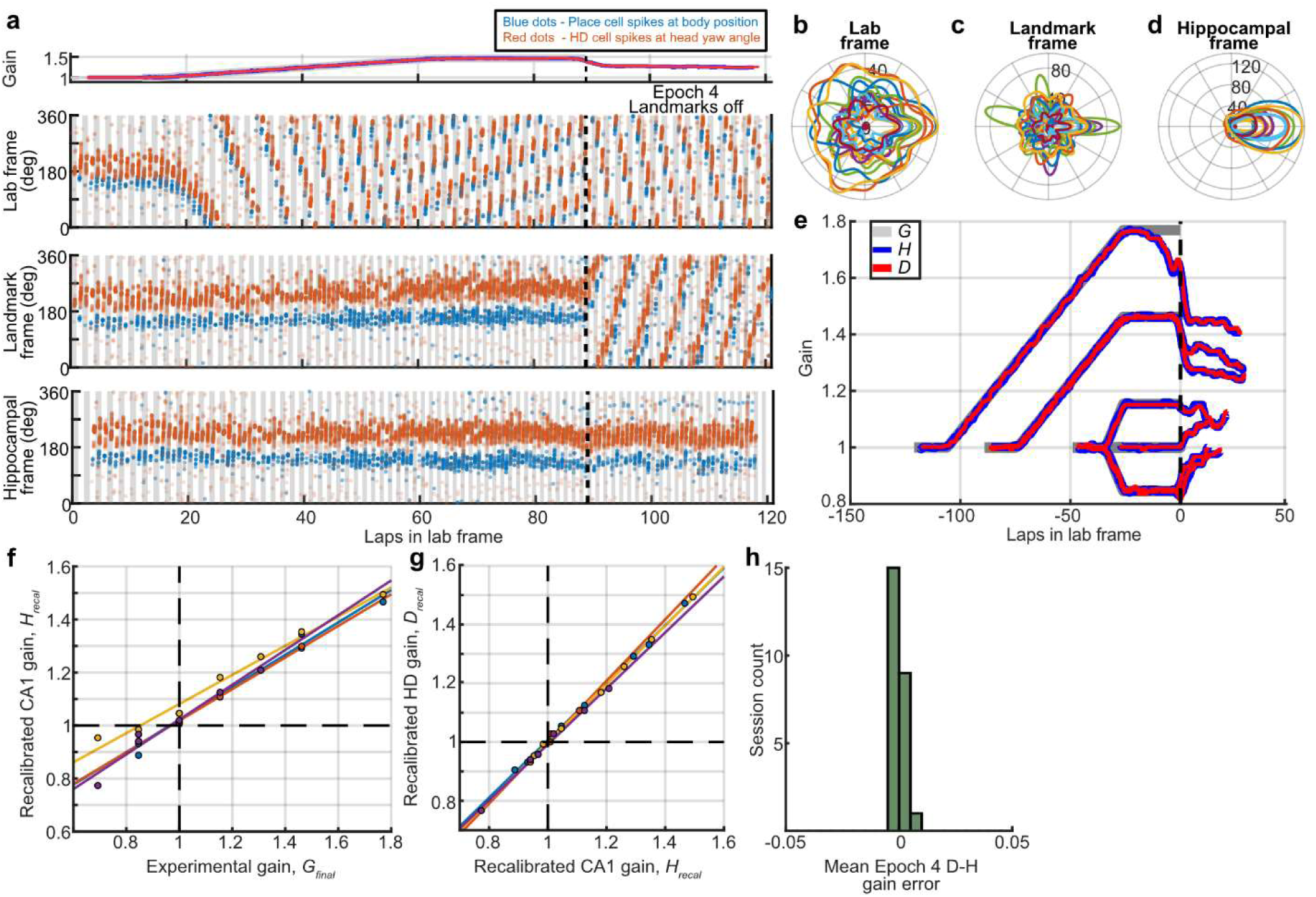
Tightly coupled recalibration of HD and CA1 path-integration gains. **(a)** Example raster plot of path integration recalibration in a place cell and a simultaneously recorded HD cell for Rat 974. Figure format is the same as Figure 2f, with the addition of data from Epoch 4 when the landmarks were extinguished (data to the right of the vertical black dashed line in all panels). The top panel shows the tight correspondence between the hippocampal population gain (blue line), the HD gain (red line), and the experimental gain (gray line) in Epochs 1-3, indicating landmark control. After landmarks were extinguished, the hippocampal gain and HD gain dropped together to 1.29, a value higher than the baseline path integration gain of 1 and recalibrated towards *G_final_*. Note that, due to the sliding window used for gain estimation, the gain traces appear to move away from *G* before landmarks turn off. As expected in a landmark-control session, the spikes display a consistent horizontal alignment relative to the landmarks in Epochs 1-3, indicating stable spatial and directional alignment relative to landmarks. In Epoch 4, the landmark frame was defined assuming the gain was *G_final_*, even though landmarks were off. Both cell types immediately started firing progressively later or earlier in the assumed landmark and lab frames, respectively, indicating a cell gain greater than the baseline gain *G* = 1. In the bottom panel, the spikes are plotted relative to the hippocampal frame (i.e., the position in the hippocampal place map as integrated from the hippocampal gain). The HD cell maintains a stable tuning in this frame indicating that the firing fields drift in tight concert in Epoch 4. **(b)** Head direction tuning curves in the laboratory frame during running for all HD cells recorded from Rat 974 in Epoch 4 across all landmark-control sessions. HD cells exhibited disrupted tuning in the lab frame. The peak firing directions of each cell are aligned to 0 degrees for visualization. **(c)** Tuning curves from the same cells as panel (b) in a fictive landmark frame calculated as if the landmarks were still present in Epoch 4 and continued to rotate at *G_final_*. HD cells exhibited stable tuning in this fictive landmark frame. The preferred firing directions of each cell are aligned to 0 degrees for visualization.

### HD gain did not exhibit recalibration during non-locomotor head scanning events

We investigated whether head direction cell recalibration was tied directly to place cell recalibration, or whether it might reflect (at least in part) a separate recalibration of angular head velocity (AHV) inputs into the head direction system. We analyzed moments when updates in position and head angle were incongruent, specifically during non-locomotor turning and head-scanning behaviors. These head-scanning behaviors were characterized by lateral head movements with minimal changes in body position. We first extracted head scans during Epoch 4 (landmarks off) based on a previously described algorithm using the animal’s head motion relative to the track (Figure 4a; Monaco et al. (57); see **Methods**). We then created the HD tuning curve for each HD unit based on spiking activity during these head scans. To decode the underlying HD gain, we applied candidate HD gain values *D* between 0.3-1.7 to the actual HD motion and obtained the putative internal HD associated with each spike (Figure 4b). Since the internal HD signal is stable in the hippocampal frame during running periods between head-scanning events, we aligned the HD of each head scan by applying the instantaneous hippocampal gain (which matches the HD gain during forward locomotion) to the animal’s body movements along the circular track; in other words, we modeled the HD gain to be equal to the hippocampal gain during running and to be one of the candidate HD gain values during head scanning. Among the candidate *D* values, the value that most closely matches the true HD gain during head scanning should produce a tuning curve most similar to the ground truth tuning estimated from head scans in Epoch 1. Hence, for each candidate HD gain value, we quantified the similarity between the resulting tuning curve and the tuning curve during head scans in Epoch 1 using circular cross-correlation; the candidate gain with the highest similarity score was assigned as the HD gain during the head-scanning events.

**Figure 4.**
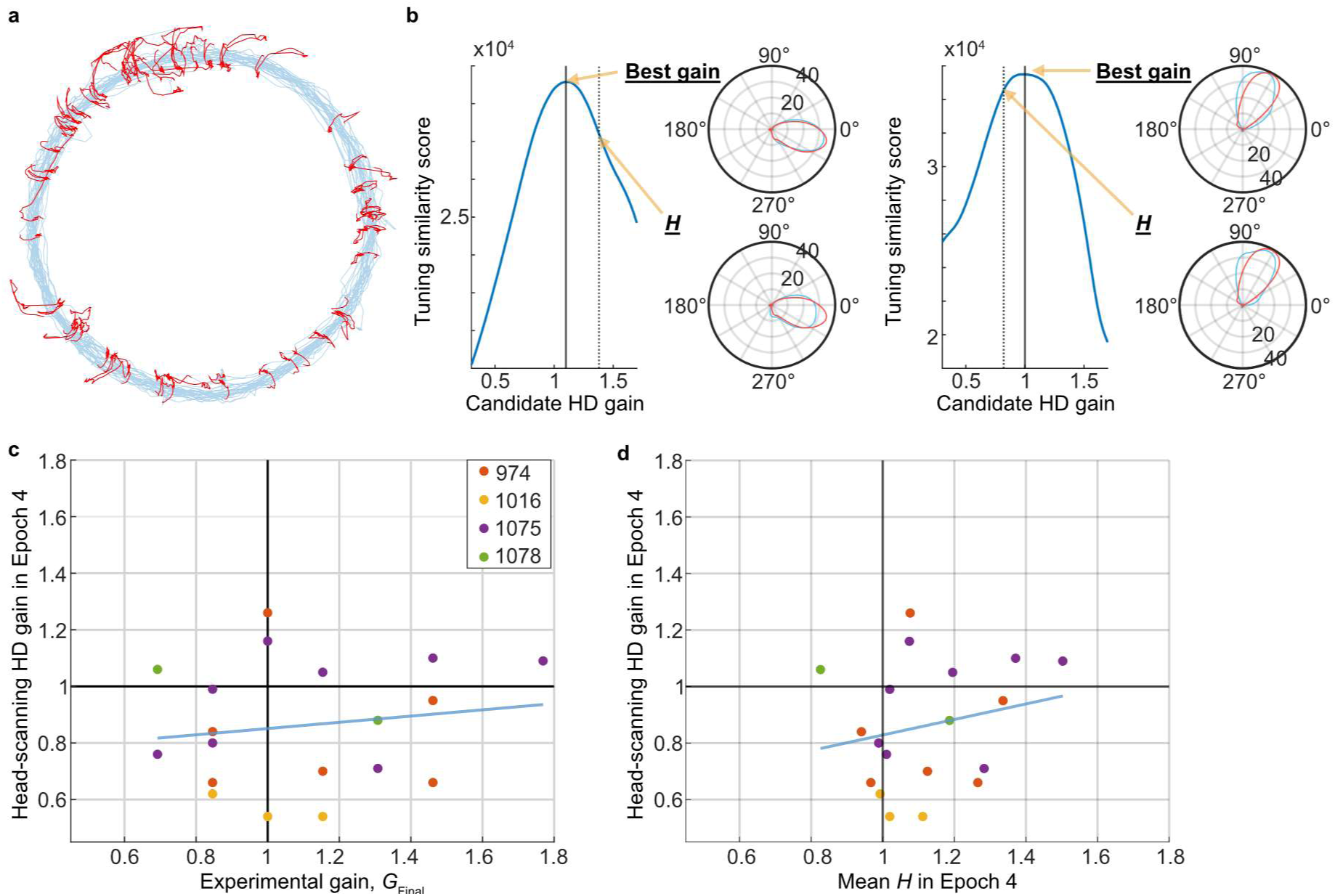
Path-integration gain recalibration occurs downstream of the HD system. **(a)** Trace of the rat’s head position during Epoch 4 from an example session for Rat 1075. The blue traces indicate when the animal was running along the circular track. The red traces indicate non-locomotor head scanning behaviors. **(b)** Schematic of the analysis to decode the HD gain during head-scanning from two representative sessions. For each unit, the tuning similarity scores are graphed as a function of candidate HD gain (left), along with polar plots of HD tuning curves for two examples (right). The top polar plot reflects the gain with the highest similarity score (producing an HD tuning curve most similar to the Epoch 1 tuning curve) and the bottom polar plot reflects the gain value of the hippocampal gain during Epoch 4. The light blue and the red lines indicate the tuning curves during Epoch 4 and Epoch 1 head scanning behaviors, respectively. **(c)** Head-scanning HD gain in Epoch 4 vs. final experimental gain. The estimated HD gain in Epoch 4 during head scanning was uncorrelated with the experimental gain in Epoch 3. Each dot corresponds to an individual session. The analysis included all sessions with HD cells from all unharnessed rats (see Methods). The light blue line is the best-fit line across the data points. Slope of recalibration: *β* = 0.206, SE = 0.310, *t*(17) = 0.665, *p* = 0.515; *t*-statistics from the linear mixed-effects model. **(d)** Head-scanning HD gain in Epoch 4 vs. mean hippocampal gain in Epoch 4 for the same sessions as in (c). The estimated HD gain in Epoch 4 during head scanning was uncorrelated with the hippocampal gain during Epoch 4. Slope of best-fit line: *β* = 0.167, SE = 0.175, *t*(17) = 0.956, *p* = 0.353.; *t*-statistics from the linear mixed-effects model.

We applied this head-scanning HD gain estimation method to our experimental data and discovered that the decoded gain did not correlate with the experimental gain in Epoch 3 (Figure 4c), indicating a lack of HD angular gain recalibration. The HD gain during head-scanning events was also uncorrelated with the corresponding hippocampal gain during Epoch 4 (Figure 4d). Together, these results indicate that the expression of path-integration-gain recalibration in HD cells is gated by behavioral state (forward locomotion vs. non-locomotor head-scanning) and/or feedback from place cells.

### HD cells and place cells exhibit a slow, biased, and partially incoherent shift relative to landmarks in landmark-control sessions

Given that HD recalibration was not directly tied to place cell recalibration in Epoch 4, we investigated whether Epochs 2-3, when the path-integration gain was undergoing recalibration, showed signs of dissociation between the place cell and the HD systems at a finer scale despite the overall coherence in gain. Our prior study showed that, in landmark-control sessions, place fields exhibited a slow, systematic shift relative to the visual landmarks across laps, suggesting a persistent contribution of path integration to place field location (21,58). Presumably, when *G* > 1, the visual input signals greater progress than does the incompletely recalibrated path integrator, and this influence of path integration causes place cells to fire slightly later relative to landmarks. Conversely, place cells fire progressively earlier when *G* < 1. We found that HD cells exhibited a similar shift, mirroring place cell behavior. In sessions with *G* > 1, *both* place cells and HD cells fired at progressively later locations relative to the landmarks (Figure 5a-b), while in sessions with *G* < 1, they fired at progressively earlier locations (Figure 5c).

**Figure 5.**
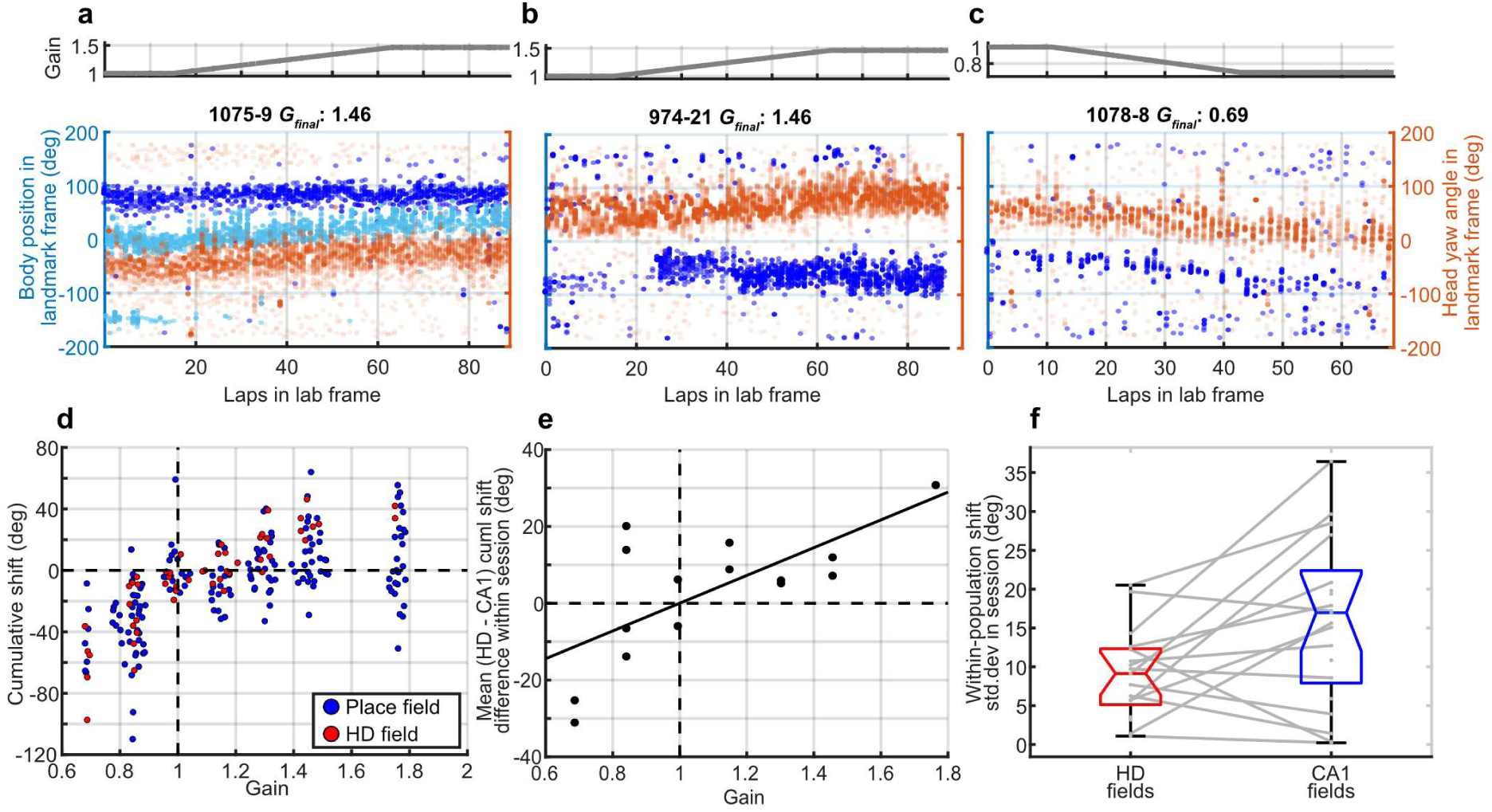
Slow, biased shift of CA1 place cells and HD cells relative to stationary landmarks during landmark control. (a) Example raster plot showing the shift of CA1 place cells (blue and cyan) and a simultaneously recorded HD cell (red) relative to landmarks during a landmark control session with G > 1. Format similar to Figure 2d. A constant offset has been added to the place and HD fields to visualize them closer together to better discern the relative shifts. The gradual lap-by-lap shift of spike positions in the place field and HD field indicates a slow shift relative to the landmark frame. In this example, the cyan place cell field shifts along with the HD field but the other field (blue) remained stable relative to landmarks. (b) Second example session with G > 1; same format. The place field (blue) diverges from the HD field and seems to have periods of forward shift, punctuated by resets back to the initial position in the landmark frame. (c) Example session with G < 1; same format. Here, the place field (blue) and HD cell (red) firing fields shift coherently in the opposite direction relative to the landmarks compared to (a) and (b), in which G > 1. (d) Scatter plot showing the cumulative shift of individual CA1 place cells (blue) and HD cells (red) as a function of the experimental gain. Positive y-axis values indicate a shift in the direction of the animal’s movement (counterclockwise) and negative values indicate a shift in the opposite direction. The place fields and HD fields show an experimental gain-dependent, shift-direction bias, indicative of conflict between the landmarks and path integration, where the path-integration gain lags behind the experimental gain. (e) Scatter plot of the within-session difference between the mean cumulative shift of HD cells versus that of simultaneously recorded place cells. HD fields shifted significantly more relative to the landmarks when compared with simultaneously recorded place fields (Difference in mean HD vs place cell shift as a function of gain with rat identity as a random effect: β = 36.1, SE = 9.90, t(15) = 3.65, p =0.00294; t-statistics from the linear mixed-effects model.) (f) Boxplot showing the distribution of within-session standard deviation of shift of HD fields and of place fields. HD fields show a smaller mean standard deviation than place fields (t(14) =-2.19, p = 0.0464), indicating a more coherent shift of HD cell tuning curves compared to CA1 place fields.

The overall trend for both cell types replicated the previously reported bias (Figure 5d), although the CA1 place cells shifted less than the HD fields on average (Figure 5e). HD cells recorded simultaneously in a session tended to shift together, exhibiting a low variance, indicating a coherent shift as an ensemble. CA1 place fields shifted with significantly larger variance, suggesting that, like the example in Figure 5a, individual place fields may weigh path integration and landmark cues differently (Figure 5f). This consistent, biased shift in both place cells and HD cells, even under strong landmark control, indicates a continuous and dynamic integration of idiothetic and allothetic information.

## Discussion

The brain seamlessly integrates internal, self-motion-based estimates of movement (path integration) with external sensory cues to support accurate navigation(32,59–64). A key variable to this computation is the path-integration gain, which associates the magnitude of displacement in the world to a distance metric in the internal, cognitive map of the animal. Given the recalibration of path-integration gain observed in hippocampal place cells (21,22), we investigated how such plasticity manifests in the head direction (HD) system. HD cells are thought to provide the directional signal that orients the cognitive map relative to the external world(31,65,66). Since rats ran unidirectionally in a circle in these experiments, coupling head direction with body position on the track, it is conceivable that the path-integration gain recalibration observed in place cells was a downstream reflection of plasticity mechanisms that occurred upstream in the circuits that integrate angular head velocity (AHV) to produce directionally tuned HD cells. Simultaneous recordings of the two cell types revealed a high degree of coupling, with HD cells exhibiting functionally identical path-integration gain recalibration with place cells during locomotion. Thus, the present experiments demonstrate that path-integration gain recalibration is reflected across the larger navigational circuit in the brain and not confined to place cells. However, the examination of non-locomotor periods (i.e., head-scanning behaviors, in which the animal changes its head direction with no concomitant changes in body location) showed no evidence of gain recalibration in the HD cells, suggesting that the plasticity resulting in the path-integration gain recalibration observed during locomotion cannot be solely driven by plasticity of AHV inputs in the head direction network.

The observed coupling in recalibration between place cells and HD cells could result from plasticity occurring either 1) in the AHV inputs onto the HD system, which entrains the hippocampal map (feedforward model; (32,67,68) or 2) downstream of the HD system in the place or MEC grid-cell system, which then entrains the HD activity (feedback model; (10,36,69)). The feedforward model predicts that the bump of activity on the putative HD ring would update during head scans according to the recalibrated path-integration gain, whereas the feedback model predicts that the HD bump would update with a baseline gain given the absence of the corresponding positional update signal provided from the place cells. Thus, the results of this study appear to support the feedback model. This interpretation is reasonable, especially given the low angular speeds that occurred (typically less than 60°/sec) as the animal moved forward along the circular track when the experimental gain of the landmarks was altered. There are alternative explanations that cannot be discounted, however. We consider two plausible alternative explanations here (although there may be others). First, it is possible that path-integration recalibration in the HD system (or elsewhere) depends on behavioral context and that the HD system can support two distinct gains. In our experiments, landmarks did not move during head scanning behaviors, providing visual feedback that *G* = 1 during head scans. Moreover, proprioceptive signals of head direction relative to the body (such as neck angle signals), fixed to the non-moving landmark frame via the feet during non-locomotor head-scans, might have served as a powerful anchor to reinforce an angular gain of *D* = 1. The carriage following the animal also provided an anchor for how much the head has turned (e.g., when the animal stopped and looked left relative to its running direction, it always saw the left wall of the carriage, and vice versa). Thus, the combination of visual and proprioceptive signals coherently indicating *G* = 1 during non-locomotor head scans, on the one hand, and a manipulated AHV gain during forward locomotion (with minimal head movement), could provide sufficient information to support distinct AHV gain calibration for these two distinct behavioral regimes (forward locomotion vs. non-locomotor head scans). Second, it may be that the manipulation in Epochs 1-3 (landmarks on) indeed recalibrated a single, coherent AHV path integration gain, but that “foot-anchored” proprioceptive cues and visual feedback from the cart (which was still visible during Epoch 4) provided a stable *D* = 1 signal when the landmarks were off that overrode the recalibrated AHV path integration signal onto the HD cells, analogous to the visual landmarks overriding path integration gain during Epochs 2 and 3. Irrespective of these alternatives, the maintenance of the relative firing positions between HD cells and place cells for dozens of laps underscores the crucial role of reciprocal interactions between these systems, as expected from previous findings (31,39).

Although the current study shows that the path-integration gain recalibration observed in place cells can occur without an accompanying recalibration in the AHV input during non-locomotor behavior (with the caveats of potential behavioral-context-dependence in mind), it is unknown whether the recalibration of the AHV input onto the HD ring can occur independently of observing path-integration gain recalibration in place cells. AHV gain recalibration does occur in the vestibulo-ocular reflex (70,71), and analogous recalibration of the HD system could be tested in our experimental apparatus by placing the animal in the center of the Dome and rotating its head in place as gain is applied to the visual scene as a function of the momentary AHV. If this manipulation were to induce AHV recalibration onto the HD cells, place cell activity could then be measured by letting the animal locomote along the circular track at the periphery of the Dome. We predict that such a manipulation would cause place fields to show recalibration (although the same caveats about behavioral-context-dependence would hold for this experiment).

Ajabi et al. (72) reported that HD cells in the ADN of mice recalibrated in the same direction as the constant-speed rotation of a visual cue in a prior light session. The mice were confined to a small platform in the center of the environment, but these authors did not record place cells in this experiment, making it unclear how place fields were affected. Moreover, because the direction of the cue rotation was decoupled from the animal’s head motion, it is unclear whether the recalibration reported in their paper reflects changes in the path-integration gain *per se* or an additive bias in HD estimation. That is, the plasticity demonstrated by Ajabi et al. may reflect, at least in part, changes in the balance of the inputs that maintain stability of the HD “activity bump” in the absence of movement, resulting in an experience-dependent, clockwise or counter-clockwise drift of the activity bump regardless of the momentary AHV. Ajabi et al. suggested that the recalibration of HD cells seen in their experiment might explain (via a feedforward model) the place field recalibration of Jayakumar et al. (21), but the lack of recalibration of HD cells during head scanning in the present study demonstrates that a more complex phenomenon is at play.

The present results demonstrates that under a closed-loop cue rotation dependent on the animal’s momentary velocity, the path-integration gain of HD cells in Epoch 4 was directly proportional to the previously experienced visual gain (as we had previously shown for place cells; (21,22)). Moreover, the coherence of the place and HD systems was maintained across various conditions (landmark control, landmark failure, and recalibration in the absence of landmarks) under which the dominant influence ranged from visual landmarks to self-motion cues. The landmark failure conditions are especially noteworthy. While a previous study has reported a session where the two systems remained coupled upon sudden unmooring of the internal map from the external world, the internal representation re-anchored to the external world after a brief interval (31). Our results demonstrate that even under a prolonged duration of such unmooring, sustained for tens of minutes of conflict between self-motion and landmark cues, the two systems remain tightly coupled. This result suggests that the coupling between place cells and HD cells is not driven by a common input from a specific modality onto the network; rather, the coupling arises from intrinsic functional connectivity between the two networks. The coupling is likely implemented by the bidirectional anatomical connections between the regions, where thalamic HD cells send polysynaptic input to the hippocampus via the parahippocampal regions (68) and the retrosplenial cortex (67), while the hippocampus sends a feedback signal to the thalamus via the dorsal subiculum (10,36). The binding of the coupled population to external visual landmarks may be provided by the activity of functional cell types in the retrosplenial cortex that mediate between visual cue and head direction signals (73,74). Together, a tightly coupled mechanism governs the response of place cells and HD cells to conflicts between internal (path integration-based) and external (visual) cues.

Although the place cells and HD cells were tightly coupled on average in their responses to the manipulations in this experiment, we discovered a degree of incongruence at a more granular scale. Extending the previous observation in the hippocampus (21), the fields of both place cells and HD cells exhibited a slow, biased shift relative to visual landmarks when the landmarks were moving in Epochs 2 and 3. The control of the direction of the shift by the visual gain suggests that the shift is a manifestation of the impact of self-motion cues counteracting the much stronger influence of the landmarks. The existence of field shifts in HD cells is in agreement with previous studies indicating that the HD system weighs visual cues and path integration cues (39,75) and observations of remapping of HD preferred firing directions depending on the magnitude of path-integration error (76). Despite the existence of shifts in both populations, CA1 place cells exhibited a more heterogeneous shift than HD cells, suggesting that different CA1 place cells may weigh path integration and landmarks differently. Because CA1 lacks a strong recurrent collateral system, individual CA1 cells are more likely to respond heterogeneously to differential inputs compared to regions like CA3 (77–80) or putative ring attractor circuits, which are the consensus model underlying the HD system (81–85). Differential cue weighting has been previously reported when distal landmarks on the wall and proximal cues on the track were rotated against each other. In these experiments, HD cells were mostly controlled by distal cues (47), while CA1 place cells showed a heterogenous locking to the two sets of cues and CA3 place cells were locked more strongly to proximal than distal cues (77). Similarly, CA1 place cells showed a heterogeneous display of backward shifts of their place fields during unidirectional movement, whereas HD cells were more homogeneous and showed different patterns of backward shifts across experience and novelty compared to CA1 and CA3 (42,86); see also (44). Thus, each component of the navigation system, despite being coupled, nonetheless is influenced by different cues provided by distinct anatomical inputs to represent direction and location. The observed heterogeneity in CA1 may also reflect the distinct properties of CA1 cells between the superficial and deep layers (87) and along the proximodistal axis (88), but we lacked the resolution and sampling density to test these hypotheses in the current experiment. Lastly, the different cue weighting between the place cells and HD cells poses a question about how cells in the hippocampal formation with conjunctive coding of place and HD respond to cue conflict. For instance, MEC contains grid x HD cells (89), while LEC contains cells that encode the animal’s egocentric HD relative to external items (90). Hippocampal place cells also exhibit HD modulation, under some conditions and to varying degrees (91–93). It is possible that these conjunctive cells exhibit a higher weighting of self-motion cues compared to nonconjunctive spatial cells, due to the HD input influencing their firing; thus, the conjunctive cells would constitute a spatial map that shifts coherently with the internal directional signal.

An unexplored question is whether the different HD circuits exhibit the properties described here. While the HD cells were recorded from the thalamus in this study, they are found across the brain including the presubiculum (24,94), posterior parietal cortex (95), postrhinal cortex (96), retrosplenial cortex (95), and dorsal striatum (97). Although the cells from the anterior thalamus in the present study showed the same amount of path-integration gain recalibration (Figure 3g**, S1e**), other components of the brain-wide HD system might show distinct profiles of gain recalibration. Moreover, recording from AHV cells that provide the angular input to the HD ring, such as units in the dorsal tegmental nucleus and lateral mammillary nucleus (26) and the retrosplenial cortex (29,56), would provide important information about whether the tuning curves of these cells can be altered with experience, or whether they remain fixed and any recalibration of the HD circuit is downstream of the sensory inputs (see Secer et al. (58) for a computational investigation of necessary conditions and potential sites of plasticity of path-integration gain recalibration in a ring attractor network).

Our study demonstrates the coordination between place cells and HD cells that maintains a coherent spatial map. Path-integration gain recalibration is likely a distributed process involving a tightly coupled mechanism between the hippocampus and the HD system, which may occur independently of the AHV input onto the HD ring. The findings indicate that the brain can maintain a self-consistent spatial representation even during the plastic process of path-integration gain recalibration to preserve path integration, a computation that is key to navigation regardless of the presence of landmarks (9,12,16,32). Furthermore, the profiles of the gradual shift of the fields of the two populations suggest that, even under a shared gain, the exact dynamics of the two networks differ in reliance on distinct spatial cues, imposing a limit to the degree of fine-scale coupling and the information used to compute these spatial metrics.

## Methods

### Subjects

Four Long–Evans rats (Envigo Harlan; 1 males [number 974] and 3 females [numbers 1016, 1075 and 1078]) were housed individually on a 12:12 h light:dark cycle. All training and experiments were conducted during the dark portion of the cycle. The rats were 5–8 months old and weighed 250–550 g at the time of surgery. Data from individual animals are shown in each figure to allow visual comparison between male and female rats. As no obvious differences were observed between sexes, data from all rats were combined for statistical analysis. All animal care and housing procedures complied with National Institutes of Health guidelines and followed protocols approved by the Institutional Animal Care and Use Committee at Johns Hopkins University.

### Dome apparatus

The Dome (50) is a planetarium-like virtual-reality apparatus with a hemispherical fibreglass shell (2.3-m inner diameter). Rats locomoted near the outer periphery of an annular table (152.4-cm outer diameter, 45.7-cm inner diameter) centered within the Dome. Visual landmarks were projected onto the inner spherical surface of the shell (Figure 1a). In addition, an overhead circular band, non-overlapping with the landmarks, was projected throughout the experimental session to provide circularly symmetric illumination inside the Dome throughout the experiment (even when landmarks were not present in Epoch 4). A near-infrared camera provided an overhead view of the experiment. The 3D position and orientation of the head of the rat were estimated in real-time (45 fps) using a specialized 3D head tracking system (84); the system uses a single camera, specialized software, and a set of markers mounted on the rat’s neural recording implant in a known 3D geometry.

An enclosure, attached to a rotating pillar at the table center via radial boom arms, covered a 45° region around the rat. The enclosure kept the rat at the outer periphery of the table and carried the transmitter unit of the neural recording system (see **Neural recording**). The rat would run approximately at the radial center of the enclosure. This was about 10 cm from the edge of the table, making the lap circumference approximately equal to 4.2m. The motorized central pillar moved the enclosure along with the rat’s head position, such that the rat was kept near the center of the enclosure. The motor was actuated only when the rat moved counterclockwise to encourage unidirectional running; the barrier at the back of the enclosure prevented the rat from running clockwise past the enclosure. Liquid reward (50% diluted Ensure®) was dispensed on the table through a feed tube on a boom arm attached to the central pillar diametrically opposed to the enclosure; the amount and timing of reward was controlled by a micro-peristaltic pump on the central pillar. This boom arm also carried a plastic spreader and paper towels that wiped up or spread out the scent of urine and uneaten food, as well as pushed feces off the table, reducing the salience and stability of local olfactory cues. All the nonprojected cues available to the rat were either circularly symmetric (nonpolarizing) or moved along with the rat.

### Training

Over 2–3 days, we familiarized the rats to human contact. The rats were placed on a controlled feeding schedule to reduce their weights to approximately 80% of their ad libitum weight, whereupon they were trained to run for a liquid reward on a training table in a different room from the experimental room. The training table had the same dimensions as the table in the Dome, but with no enclosure or other automated systems. Reward droplets were manually placed at arbitrary locations on the track in the path of the running rat, and the experimenter attempted to lengthen the average interval between rewards to maintain behavior while delaying satiation. Training continued until the rats consistently ran at least 40 laps (often up to 100 laps) without intervention or encouragement from the experimenters, usually taking 2–3 weeks.

### Electrode implantation

After training, rats were implanted with hyperdrives containing 18 independently drivable nichrome tetrodes (diameter of wire = 0.0178 mm, Kanthal, Sweden), two of which served as references. The hyperdrives were fabricated in the laboratory using an in-house design with a 72-channel interface board (EIB-72-QC, NeuraLynx, Bozeman, MT, USA). The tetrodes were grouped into two bundles to simultaneously target CA1 and regions known to have HD cells, primarily the ADN of the thalamus. Prior to surgery, the tetrode tips were gold-plated to an impedance of ∼150 kΩ using a nanoZ electroplating system (White Matter LLC, Seattle, WA, USA). The implantation coordinates were determined during surgery based on the location of the septal pole of the hippocampus, which is situated directly above the ADN. Briefly, a recording electrode was inserted in a grid search around coordinates ML 1.4 mm and AP-1 mm relative to bregma to locate the hippocampal septal pole. Once the septal pole was found, the hyperdrive was implanted such that the ADN bundle of tetrodes was placed over the septal pole. Given the geometry of the hyperdrive and orientation of implantation, the center of the CA1 tetrode bundle was approximately at coordinates ML 2.4 mm and AP-3.3 mm relative to bregma. Following surgery, 30 mg of tetracycline and 0.15 ml of a 22.7% solution of the antibiotic enrofloxacin were administered orally to the rats each day. After at least four days of recovery, we began slowly advancing the tetrodes; we resumed food restriction and training within seven days of surgery.

### Electrode adjustment

Tetrodes in the CA1 bundle were advanced less than 40 µm per day once they were close to CA1. The location of each tetrode relative to the CA1 pyramidal cell layer was judged using the polarity of sharp waves and intensity of ripples in the electroencephalogram (EEG) signal captured on one electrode of each tetrode, as per well-established procedures. Tetrodes were judged to be correctly placed when ripples were intense and multiple units were visible on the pairwise electrode projections of spike amplitudes. Tetrodes targeting the ADN were initially advanced quickly, using characteristic electrophysiological patterns of the cortex, corpus callosum, hippocampal septal pole and fimbria of the hippocampus as landmarks. The advancement of the tetrodes was slowed upon reaching the quiet zone of the ventricular system above the ADN. Tetrodes were judged to be correctly placed when they reached a region with head direction cell activity, identified by direction-specific firing as the rat was passively rotated around the yaw axis while resting on a turntable. In a small subset of sessions, we recorded HD units that were encountered during tetrode advancement in the cingulate bundle above the hippocampus.

### Post-surgery training

During the days of tetrode advancement, we food-restricted the rats. On (typically) day 3 of food restriction, we introduced them into the Dome with stationary visual landmarks. The process typically started with a free exploration of the tabletop for 1–2 hours (broken up into half-hour sessions). After this initial familiarization session, rats were encouraged to run along the periphery of the table counterclockwise in pursuit of liquid reward. In the following two days, animals were trained to run counterclockwise around the periphery of the Dome table in pursuit of reward with the neural recording cables and position-tracking markers attached to the hyperdrive on their heads. Reward was dispensed at random spatial intervals sampled from a uniform distribution, whose range depended on the rat’s performance. The range was gradually increased to maintain running performance until the pre-surgery performance criterion was reached (typically 7–10 days).

### Neural recording

Experimental sessions began once the tetrodes were judged to be in CA1, at least one HD cell was recorded, and the rat was running at least 40 laps inside the Dome. During the sessions, a unity-gain neural recording headstage (HS-72-QC, NeuraLynx) was attached to the implanted hyperdrive. The signal was filtered (0.1-8000 Hz), digitized at 30 kHz, and sent to the Cheetah computer wirelessly through a transmitter unit (FreeLynx, NeuraLynx). Further filtering was performed by the Cheetah 5 recording software to extract the LFP (0.1-600 Hz) and spikes (600-6000 Hz).

### Experimental control

Three computers were used to run the experiment. Their purposes were (1) general experiment control, (2) neural recording (Cheetah computer), and (3) video tracking and recording. Multiple independent programs, called nodes, performed individual tasks and were in continuous communication with a master node running on Computer #1 and with each other through a software framework called Robot Operating System (ROS) (99). Pseudo-random digital pulses sent from Computer #1 were time-stamped and recorded as events on the Cheetah computer to enable the post hoc synchronization of the data streams recorded on the two computers. Details on the hardware and software integration and experimental control are available in (50).

### Experimental procedure and gain selection

On each experimental day, sleep sessions were recorded for 20 mins before and after the experimental session. During the sleep sessions, rats either rested on a pedestal outside of the Dome (rats 974, and 1016) or in a cylindrical enclosure inside the Dome (rats 1075 and 1078). These data were used post hoc to confirm the recording stability of single units during the experiment.

During an experiment, an array of visual landmarks was moved as a function of the rat’s motion. The motion of the visual scenes was governed by the experimental gain, *G*, such that *G* x (rat’s speed in lab frame) = (rat’s speed in landmark frame). For instance, landmarks were stationary at *G* = 1, landmarks moved proportionally to the animal’s movement in the same direction as the animal when *G* < 1, and in the opposite direction as the animal when *G* > 1. An experimental session consisted of four epochs. In Epoch 1, rats ran 15 laps with stationary visual landmarks (Figure 1c). In Epoch 2, *G* was linearly increased or decreased at a constant rate to *G_final_*. In Epoch 3, *G* was held steady at *G_final_*. Finally, in Epoch 4, visual landmarks were turned off, such that only a nonpolarizing, circular ring was projected near the top of the inner shell for illumination. The values of *G_final_* were chosen to be of the form, 1 ± n/13 with n = 2,4,6,10, resulting in gains of 0.231, 0.539, 0.692, 0.846, 1.154, 1.307, 1.462, and 1.769. These values with a prime denominator were used to minimize the frequency of alignment of the lab and landmark frames; with the chosen gain values, the position of the rat relative to the two frames coincided only once every 13 laps in Epoch 3. For the first three animals, gains were typically changed at a constant rate of 1/52 per lap, such that the length of Epoch 2 was 8, 24, and 40 laps for n = 2, 6, and 10, respectively. In these animals, a high proportion of *G* < 1 sessions showed a weaker influence of the visual landmarks on the hippocampal map, such that the place fields drifted from the landmark frame (loss of landmark control). To establish landmark control in larger gain ranges, the gain ramp rate was slowed to 1/104 per lap for the last two animals. The sessions were not randomized; the gain for each session was selected such that gains were rarely repeated in consecutive sessions, and the gain manipulation typically increased in magnitude over consecutive sessions for any given animal. This ordering aimed to maximize the range of gain values with landmark control; higher magnitudes of gain manipulation were more likely to lead to loss of landmark control, and such loss of landmark control in one session often persisted in subsequent sessions. The investigators were not blinded to allocation during experiments and outcome assessment. No statistical methods were used to predetermine sample size.

### Data analysis Spike sorting

Spikes were sorted based on their peak, valley, and energy values using a custom software program (WinClust; J.J.K.). Cluster boundaries were drawn manually on two-dimensional projections of these features from two different electrodes of a tetrode. We mostly used peak and energy as features of choice; however, other features were used when they were required to isolate clusters from one another. Clusters were assigned isolation quality scores ranging from 1 (very well isolated) to 5 (poorly isolated), agnostic to their spatial-firing properties. For units that drifted within a session, an isolation quality score of 5 was assigned unless the spike cluster, despite drifting, was able to be confidently tracked and isolated across time (in which case an isolation quality of 1-4 was assigned).

### Inclusion criteria

To be included in the quantitative analyses, sessions were required to be completed without major behavioral issues or long manual interventions during the recording. Session inclusion was determined before performing any of the statistical analyses reported here. For units recorded in the tetrodes in the hippocampus, only the units with isolation quality of 4 or above were used for single-unit analyses; all units were included for the multi-unit analyses (i.e., spectral decoding analysis) given the coherence in gain among co-recorded units (**Figure S1e**). For units recorded outside of the hippocampus (mostly anterior thalamic nuclei, **Table S1**), units classified as HD cells (see the following section) were included in the analysis.

#### Head direction cell classification

HD cells were defined as units recorded in tetrodes over stereotaxic coordinates targeting the anterior thalamic nuclei that met the following criteria. First, the unit had to have a high HD information score both during running (HD information score > 0.5; forward speed > 5.8 cm/s, which is equivalent to 5°/s along the arc of the circular track) and stationary periods (HD information score > 0.4; forward speed < 5.8 cm/s), where the information score was defined using methods equivalent to spatial information scores for place cells (100). Although the firing rates were calculated using the true distribution of directional occupancies, we made a simplifying assumption of a uniform probability of direction occupancy in the information score equation due to inhomogeneities in directional sampling. The HD information score was computed during Epoch 1, in the absence of conflicting frames of reference. Further, given that the information score is known to penalize units with non-zero baseline firing rate (101), we used the firing rate after subtracting 0.8 x (minimum firing rate); however, for single-unit analyses (i.e., head scanning analysis) in which spike contamination could potentially influence the results, we only included the units that met the information score criteria without the baseline subtraction. Second, we required that the HD tuning curves of the running and stationary periods be similar (Pearson’s correlation coefficient > 0.6). Last, we required that the unit had a peak firing rate > 5 Hz.

While running unidirectionally on the circular track, the animal’s head direction was usually aligned with the track’s tangent, creating a strong correlation between its location (place) and head direction. This coupling could cause a spatially tuned cell to be misclassified as a directionally tuned one. We thus manually verified that all units that met the initial HD tuning criteria were true HD cells by analyzing periods when the animal’s place and head direction were decorrelated. For some sessions (rats 1075 and 1078), we confirmed HD tuning using recordings of the animal exploring a small open field chamber placed inside the Dome at the end of the Dome experiments. For other sessions (rats 974 and 1016), we referenced experimenter notes taken before the Dome session, which documented HD tuning while the animal was rotated in place on a pedestal. If the relative preferred directions of the simultaneously recorded, putative HD cells in the Dome matched the preferred directions of the HD cells in the experimenter’s notes, we considered them to be verified HD cells. Finally, for any remaining units, we analyzed head-scanning epochs—periods when the stationary animal made lateral head movements (see Methods: Head-scanning analysis) that occurred outside the cell’s “spatial field” (i.e., the location on the track whose tangent was parallel to the cell’s preferred direction). If the preferred direction of the HD cell during head scanning in these locations matched the preferred HD direction of the entire session, we considered the cells to be verified HD cells.

#### Spectral Decoding

The algorithm for spectral decoding of hippocampal gain is detailed in (22). Briefly, the gain was defined as the spatial frequency of individual place fields on the circular track. For a stable spatial representation in the lab frame, a typical CA1 place cell would exhibit one firing field that repeats every lap. Hence, the spatial frequency of firing would be 1 cycle/lap. If a cell repeated its field more (or less) than once per lap, its spatial frequency would be > 1 (or < 1) cycles/lap. To compute the spatial firing frequency, the unwrapped spatial firing rate for each unit was obtained by dividing the number of spikes by the time spent in the respective bins (size = 5.8 cm), only considering running epochs (speed > 5.8 cm/s). The spatial frequency was obtained by applying the spectral decoder on the spatial firing rates with a 6-lap window. The median of these unit gains was defined as the population hippocampal gain *H*.

Head direction gain was estimated similarly to the hippocampal gain but using the unwrapped yaw angle of the head. We leveraged the ∼1:1 relationship between the animal’s head yaw angle and position during locomotion, given that the rats ran laps on a circular track. Before the gain estimation, periods where head yaw and track angles deviated significantly (circular distance > 60°) were excluded, which mostly consisted of pausing and turning behaviors. Using the remaining time points, the HD gain was decoded by the same spectral decoder as above using the yaw angle-based firing frequency of the HD cells.

### Mean gain error score

To evaluate the coherence of the recorded neural population, we quantified the deviation of the unit gain relative to the population gain. For each unit *i* and for each decoding window (6 laps), the unit gain estimate *H_i_* was compared against the population gain *H*, computed with *H_i_* excluded. The mean gain error score was defined as the session mean of |*H_i_* – *H*| (21). Intuitively, the score would be zero if all units exhibited identical gains and increase with larger incoherence in gain among the units.

### Definition of landmark-failure and landmark-control sessions

We classify a given session as being a landmark-failure session if, over the course of Epochs 1–3, the ongoing hippocampal gain estimate *H* deviated from the experimental gain *G* by more than an absolute value of 0.05 for at least one lap. This deviation indicated that the visual landmarks had, at least for a sustained period, lost their anchoring influence on the hippocampal place map. If the classification threshold wasn’t crossed, the session was classified as a landmark-control session.

### Calculating the position in the hippocampal frame of reference

The displacement of the animal in the hippocampal frame of reference is defined as follows:

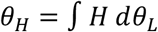

where 𝜃_H_ is the displacement in the hippocampal frame, 𝜃_L_ is the angular displacement in the lab frame, and the integration is taken with respect to the animal’s angular displacement in the lab frame, 𝑑𝜃_L_. Computationally, this position in the hippocampal frame was estimated by taking successive angular displacements of the animal in the lab frame, scaling these angular displacements by the hippocampal gain at that angle, *H*, and computing the cumulative sum of these scaled incremental displacements.

### Field detection and tracking

For landmark-control sessions, we tracked the lap-to-lap dynamics of place fields and HD cell firing fields in the landmark frame. First, occupancy-corrected firing rate was computed with respect to the unwrapped position/head yaw angle. Here, the spike count was divided by the spatial/angular occupancy, both smoothed using a Gaussian kernel of 5.8 cm bandwidth and only using running epochs (speed > 5.8 cm/s). Second, candidate fields were identified as positions/yaw angles that exhibited elevated firing rates (20 % of maximum firing). Third, candidate fields were defined as bona fide fields if their field sizes were within 10-180° and were active for at least 12 laps. Finally, for each identified field, a time-frequency ridge detection algorithm (MATLAB tfridge function) was used to track the location of maximum firing across laps. Together, the algorithm tracked the field boundaries, peak location, and peak firing rate for each traversal through the identified fields.

### Head-scanning analysis

We measured the angular path-integration gain of the HD system when rats turned around in place or performed head scans before/after reward consumption. To extract these behavioral epochs (collectively referred to as head scans hereafter), we used a modified version of a published algorithm to extract head-scanning events (57). First, we defined candidate head-scanning events by excluding time points in which the animal was running along the track. Running was defined by first selecting time points in which the animal’s velocity along the track (referred to as body speed hereafter) exceeded 5.8 cm/s and then further selecting time points where the animal’s head yaw was aligned with the track (yaw-body offset < one standard deviation of the offset distribution). The latter criterion allowed us to avoid erroneously excluding time points where a head scan along the track showed up as a body speed greater than the threshold, given that the body position is defined as the center of the head. Second, after removing these running moments, we selected time points where the animal’s yaw speed was greater than 5°/s. Third, for each chunk of time points that passed the previous criteria, 50 ms was padded to the ends, and events close to each other (≤ 0.5 s gap between consecutive events) were merged. Lastly, we applied a filter to only select chunks of putative head-scanning time points that showed prominent head scans (yaw coverage > 45°, furthest distance of head from track center > 1.5 * track half width, body coverage < 52 cm [, equivalent to 45° along the arc of the circular track]). Given that the exact behavior was different across animals and across sessions, some of the parameters were manually adjusted to improve accuracy.

Using the extracted head-scanning events in Epoch 4, we computed the HD tuning curves of each HD unit. To decode the underlying HD gain, we performed a grid search by applying a range of candidate HD gain values (0.3-1.7) to the actual HD motion, providing the putative internal HD associated with each spike. To account for the interleaving running epochs, where the internal HD representation is updated by the hippocampal gain, the HD of each head scan was aligned by applying the instantaneous hippocampal gain to the animal’s body movements along the circular track. The resulting tuning curve would be most similar to the original tuning curve measured in Epoch 1 if the applied gain was nearly identical to the ground truth HD gain. Hence, we computed the similarity score of each tuning curve relative to the HD tuning curve obtained from head-scanning behaviors in Epoch 1. Here, the similarity score was defined as the maximum value of the circular cross-correlation between the two tuning curves, where a cross-correlation was applied instead of a simple correlation to account for the accumulated drift in tuning during the gain manipulation (Epochs 2 and 3). The candidate gain value that maximized this tuning curve was defined as the head-scanning HD gain. To exclude units/sessions with erroneous estimates due to a low number of head scans or aberrant HD firing patterns, we limited the analysis to units that showed a single peak in the similarity score.

The method was verified using a simulation of a virtual rat running on a circular track at a constant speed (5 cm/s) for 5000 s. At a regular interval (every 110°), the simulated rat made a head scan (outbound head scan with coverage: 90° and speed: 20°/s). We modeled a HD cell whose activity followed a Poisson process sampled from a circular Gaussian tuning function (peak firing rate: 20 Hz, sigma: 20°). We simulated the resulting activity of the HD cell under a range of HD gains. We applied the above tuning similarity measure to the resulting spiking activity to test whether the method can correctly recover the underlying HD gain that was simulated and verified that the above method correctly decodes the underlying HD gain across a range of recalibrated gain values.

### Histology

Once experimental sessions were complete, rats were transcardially perfused with 3.7% formalin. The brain was extracted and stored in 30% sucrose-formalin solution until fully submerged. The brain was subsequently sectioned coronally at 40 µm intervals. The sections were mounted and stained with 0.1% Cresyl violet, and each section was photographed. These images were used to identify tetrode tracks based on known tetrode bundle configuration. A depth reconstruction of the tetrode track was carried out for each recording session to identify the specific areas in which the units were recorded.

### Statistics

Parametric tests were used to determine statistical significance. Pearson product-moment correlations were used to test the linear relationship between variables. Linear mixed-effects models were used to test the significance of differences in field drift between the HD cell population and the place cell population, with (Mean Drift_HD-_ Mean Drift_CA1_) as the response variable, 𝐺_final_ as a fixed-effect variable, and rat identity as a random-effect variable. Similarly, for the quantification of the head scanning analysis, the head-scanning HD gain was modeled with the experimental gain (or the mean hippocampal gain in Epoch 4) as a fixed-effect variable, and rat identity as a random-effect variable. The model was fit using the command *fitlme* in MATLAB.

## Acknowledgments

We thank the members of the Cowan and the Knierim labs for helpful comments.

## Author contributions

Conceptualization: R.P.J., J.J.K., and N.J.C.

Experiments: R.P.J., Y.S., M.F., B.Y.L., and M.S.M.

Data analysis: R.P.J., Y.S., M.F., X.C.

Supervision: J.J.K. and N.J.C.

Writing: R.P.J., Y.S., J.J.K., and N.J.C. wrote the manuscript, with inputs from all authors

## Data and materials availability

Upon publication, the data and code used in the analyses will be made available at https://doi.org/10.7281/T1DXOIB0, Johns Hopkins Research Data Repository.

---

**Figure S1:**
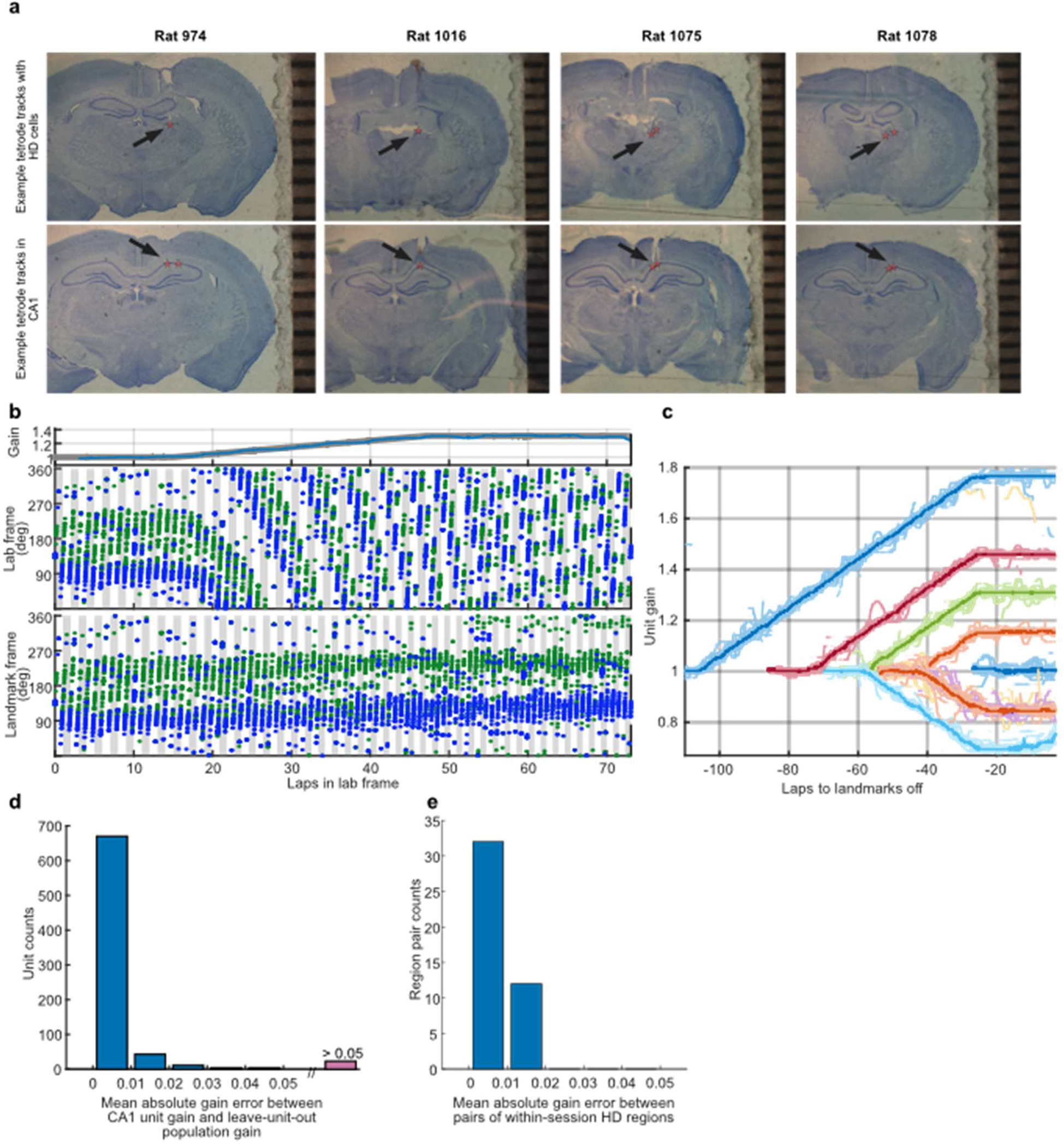
Histology and within-population coherence of hippocampal and HD units. **(a)** Example coronal slices indicating representative locations of tetrodes recording from HD cells and place cells in the five rats. Top row shows tips of tetrode tracks that recorded from HD cells (recording locations indicated by black arrow and red translucent stars; rat-wise breakdown of regions in Table S1). Bottom row shows the proximodistal location of tetrodes recording from hippocampus. **(b)** Example raster plot of two simultaneously recorded CA1 place cells from a landmark control session. The x-axis represents the cumulative distance traveled by the rat in the lab frame. *Top Panel*: Hippocampal population gain trace (blue) closely tracks the experimental gain (black). *Middle Panel*: Blue and green dots indicate spikes from two different place cells on the track relative to the lab frame (Y axis). *Bottom panel*: the location of the same spikes relative to landmark frame (Y axis). Alternating gray and white bars denote individual laps. When the landmarks began to move as the experiment gain ramped up to *G_final_* =1.31, the spatial tuning of the two place fields changed continuously in the lab frame but remained stable relative to landmarks in this session and coherent with each other. **(c)** Hippocampal unit gain traces from landmark-control sessions from one animal. Colors indicate session identity. Each light trace is an individual hippocampal unit gain; the dark traces indicate the median of the individual traces from the session. The tight coherence of gain traces throughout the course of each session indicates a coherent hippocampal ensemble. **(d)** Histogram showing the distribution across all sessions of mean absolute error between the gain trace of each hippocampal unit within a session and the population median hippocampal population gain trace over the entire session. For this analysis, the population gain trace was computed as the median of unit gains, leaving out the unit under comparison. As expected from the coherent hippocampal response seen in (c), the unit gain errors were tightly clustered near zero. **(e)** Histogram showing within-session mean gain error from pairwise comparisons between the median gain responses from HD units from one anatomical region to the median responses of units from a different simultaneously recorded region. Since the errors were tightly clustered around zero, a single HD gain was assumed in all subsequent analyses regardless of the anatomical region of origin. The gain estimate is a relatively coarse estimate, and region-dependent differences may exist with finer analyses. Given the limited number of HD cells recorded per session, claims about region-specific differences likely cannot be supported.

**Figure S2:**
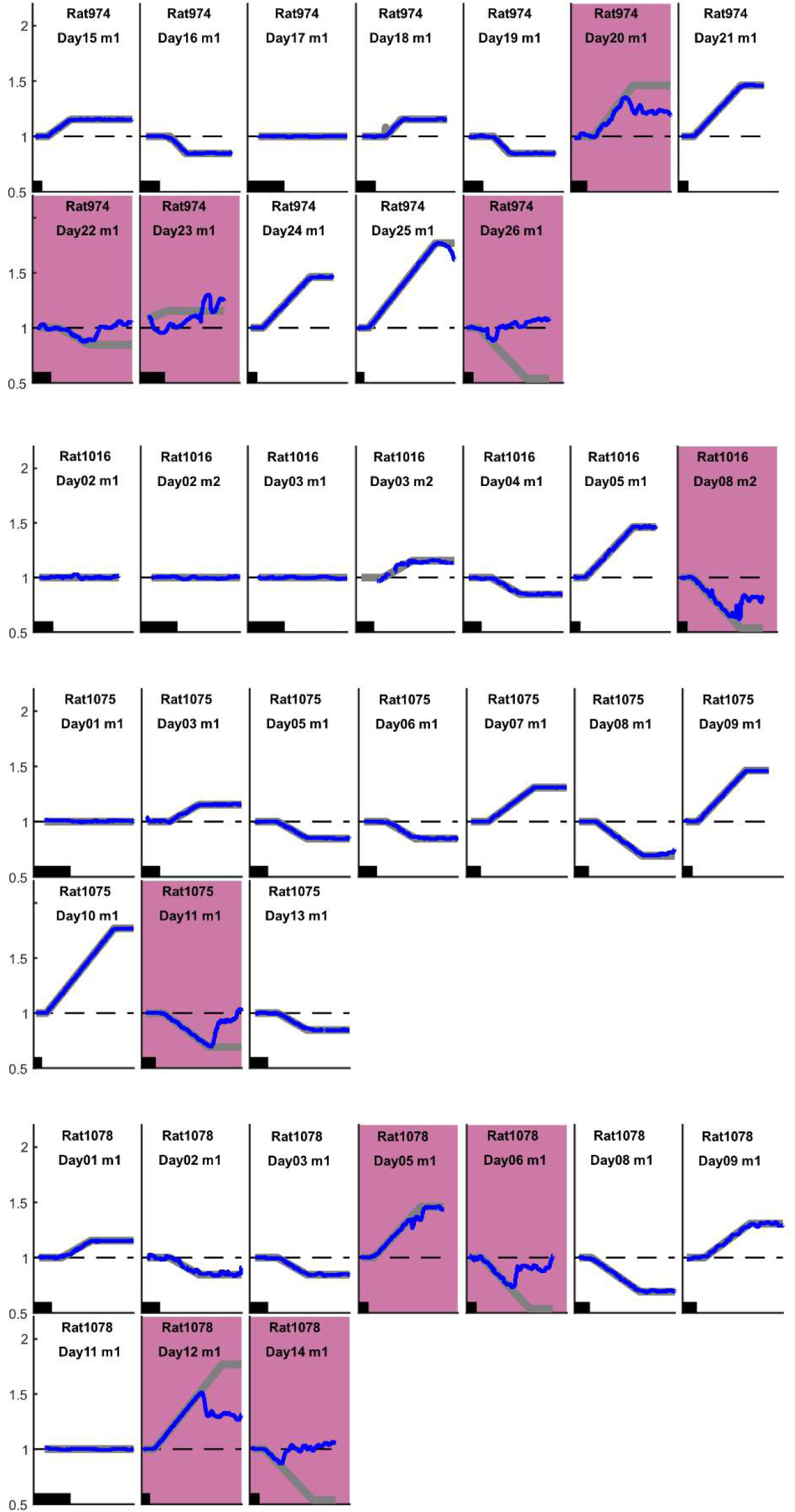
Session classification into landmark control and landmark failure. Each panel represents data from a single session during Epochs 1–3 (landmarks on). Session ID, with information about rat of origin, is denoted by the text at the top of each panel. The X axis represents the cumulative distance traveled by the rat in the lab frame. The black scale bar in each panel indicates 10 laps. The applied experimental gain *G* (gray) is plotted with the hippocampal gain estimate *H* (blue). Pink panels indicate sessions with loss of landmark control (see Methods). In the other plots, the gray and blue curves overlap, indicating control of landmarks over the place fields.

**Figure S3:**
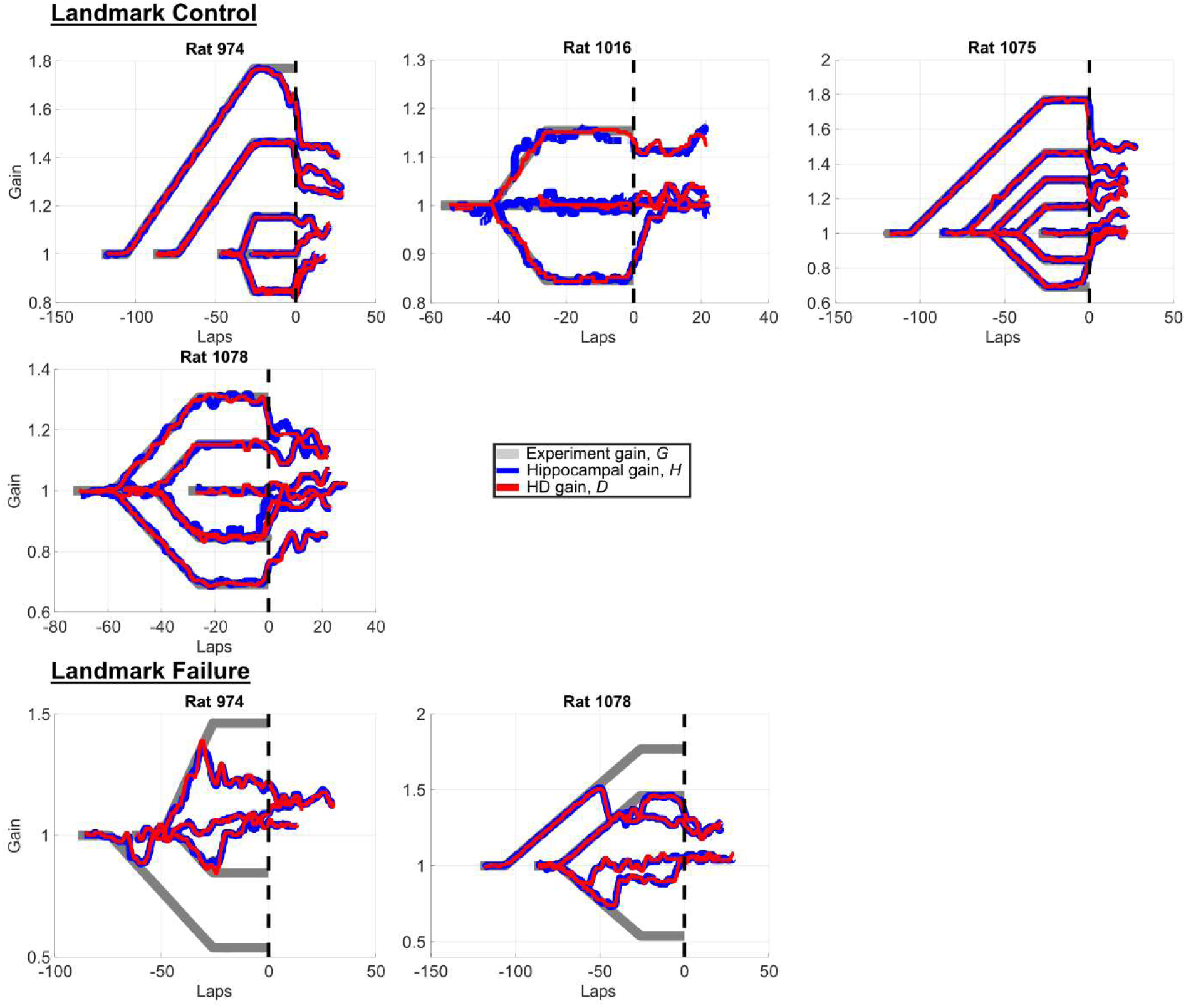
Hippocampal gain and HD gain dynamics co-evolve under all conditions Each panel shows data from sessions (Top set: landmark control, Bottom set: landmark failure) from an individual rat. X axis shows the cumulative distance in lab frame until the landmarks are extinguished (lap 0, vertical dashed line). Hippocampal population gain (blue) and the HD gain (red) evolve in tight concert with each other, regardless of whether they remain anchored by the visual cues (experiment gain, gray).

**Figure S4:**
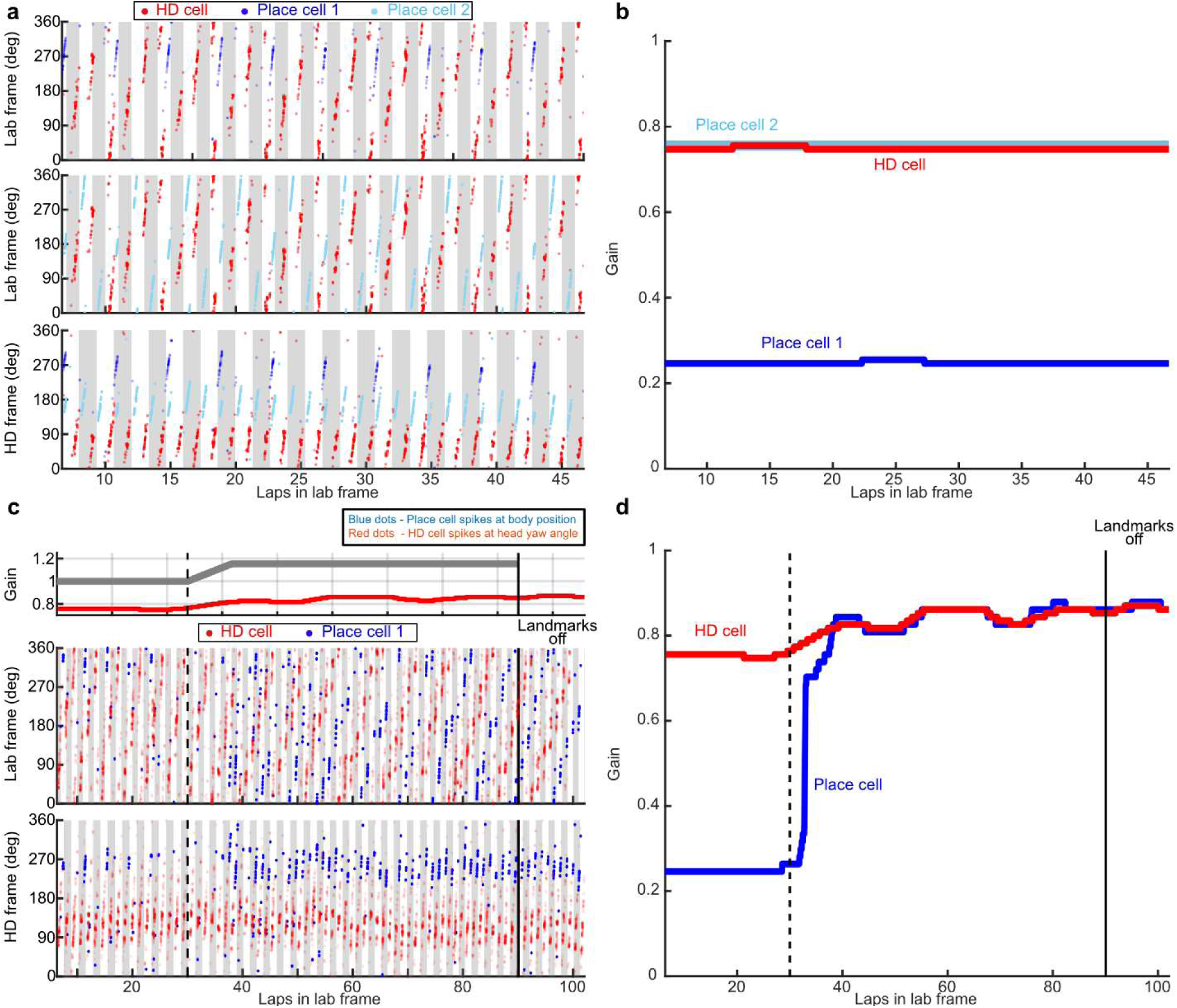
Two anecdotal sessions that violated the strict coupling typically observed between place cells and HD cells (not included in other analyses). **(a)** Spike raster plot of two simultaneously recorded CA1 place cells (blue and cyan) and an HD cell (red) during a *G = 1* session for Rat 974. Format is the same as Figure 2a, where the x-axis represents the cumulative distance traveled by the rat in the lab frame and gray and white bars indicate laps in the frame of reference indicated on the y axis. The top two panels show the spikes of the same HD cell plotted in the lab frame along with each of the place cells individually for visual comparison. At first glance, it appears as if the HD cell has three distinct stable fields in the lab frame, separated by 120°. However, the three “fields” fire on sequential laps and come back to the first firing location every fourth lap. This pattern is due to the HD cell updating with a gain of ¾ which results in this stable repeating pattern with reference to the lab/landmark external frame. Place cell 1 (blue, top panel) had one distinct stable field in the lab/landmark frame that repeated consistently once every four laps, corresponding to a cell gain of ¼. Place cell 2 (cyan, middle panel) drifted along with the HD cell and displayed a similar pattern of three fields every four laps. The firing of Place cell 1 appears to encode the *conjunction* of the HD cells with the external reference frame as the HD and Place cell 2 fields realigned with the lab/landmark frame every four laps. To visualize the two place cells’ patterns of firing, the bottom panel plots the same place cell spikes relative to the frame of reference of the drifting HD cell (computed by integrating its gain trace, similar to estimation of the hippocampal frame). The alternating gray and white bars are wider in the bottom panel compared to the top 2 panels and represent a full turn in the HD space. In this HD frame of reference, the HD cell (red) and Place cell 2 fired once every turn, while Place cell 1 fired once every 3 turns, corresponding to the periodic alignment of the HD frame and lab/landmark frame. **(b)** Gain traces of cells in (a). X-axis represents the cumulative distance traveled by the rat in the lab frame. While the gain of the HD cell (red line) stayed at ¾, Place cell 1 blue line) exhibited a gain of ¼. Place cell 2 (cyan line, occluded by HD cell gain trace) exhibited a gain similar to the HD. **(c)** Spike raster plot of an HD cell (red) and a simultaneously recorded place cell (blue) from a subsequent session. The format is similar to (a) with the addition of visualization of gain traces in the top panel since the experiment gain *G* (gray) was manipulated. During Epoch 1 (G = 1), the HD cell displayed the same drift rate as in (a) of repeating three firing fields every four laps in the landmark frame, and the place cell fired once every four laps (middle panel), corresponding to once every three turns in the HD frame (bottom panel). However, as *G* was increased from 1 in Epoch 2 (black dashed line), the conjunctive firing pattern of this cell collapsed to start drifting along with the HD cell. When landmarks were extinguished (solid black line), the place cell continued drifting coherently with the HD cell. **(d)** Gain traces of cells in (c). During Epoch 1, the Place cell (blue line) behaved similarly to Place cell 1 in the session shown in (b), exhibiting a gain of ¼. Once the gain manipulation started, however, the gain of the Place cell stopped encoding the conjunction of the HD frame and lab/landmark frame and eventually aligned with that of the HD cell.

**Table S1.** Anatomical locations of tetrode tracks recording head direction (HD) cells. The table provides a summary of the specific brain region for each recording track, organized by individual rat. Abbreviations: ADN, Anterodorsal thalamic nucleus; AVN, Anteroventral thalamic nucleus; LDN, Laterodorsal thalamic nucleus.

| **Rat Number** | **HD Region** |
| --- | --- |
| 974 | AND/LDN/Cingulum |
| 1016 | ADN |
| 1075 | ADN/AVN |
| 1078 | ADN/AVN |

